# Transcriptional profiling of sequentially generated septal neuron fates

**DOI:** 10.1101/2021.06.29.450296

**Authors:** Miguel Turrero García, Sarah K Stegmann, Tiara E Lacey, Christopher M Reid, Sinisa Hrvatin, Caleb Weinreb, Manal A Adam, M Aurel Nagy, Corey C Harwell

## Abstract

The septum is a ventral forebrain structure known to regulate innate behaviors. During embryonic development, septal neurons are produced in multiple proliferative areas from neural progenitors following transcriptional programs that are still largely unknown. Here, we use a combination of single cell RNA sequencing, histology and genetic models to address how septal neuron diversity is established during neurogenesis. We find that the transcriptional profiles of septal progenitors change along neurogenesis, coinciding with the generation of distinct neuron types. We characterize the septal eminence, a spatially distinct and transient proliferative zone composed of progenitors with distinctive molecular profiles, proliferative capacity and fate potential compared to the rostral septal progenitor zone. We show that *Nkx2.1*-expressing septal eminence progenitors give rise to neurons belonging to at least three morphological classes, born in temporal cohorts that are distributed across different septal nuclei in a sequential fountain-like pattern. Our study provides insight into the molecular programs that control the sequential production of different neuronal types in the septum, a structure with important roles in regulating mood and motivation.

## INTRODUCTION

A central question in developmental neurobiology is how a pool of seemingly uniform embryonic neural stem cells can generate the immense diversity of neuronal subtypes in the mature brain. During mammalian brain development, neural progenitors acquire distinctive spatial identities and progress through a series of temporal competence states to give rise to their neuronal progeny. There has been significant progress in defining the key molecular differences between spatially segregated progenitor domains. For example, the transcription factors EMX1 and DLX2 define the pallial and subpallial progenitor domains responsible for the production of glutamatergic and GABAergic forebrain neurons, respectively (Puelles et al., 2000). Temporal programming of forebrain progenitors is perhaps best exemplified in the developing cerebral cortex, where neurons are sequentially produced to populate the six cortical layers in an inside-out pattern (Lodato and Arlotta, 2015). One of the major challenges in the field has been to identify the molecular programs controlling the timely generation of diverse neuronal subtypes from each progenitor region within the forebrain (Kohwi and Doe, 2013). Recent advances in single-cell RNA Sequencing (scRNA-Seq) techniques have greatly increased our understanding of the molecular identity of numerous mature neuronal types (Yuste et al., 2020), and began to address the dynamic transcriptional changes underlying their specification (Loo et al., 2019). In the ventral forebrain, several studies have used scRNA-Seq to address how and when GABAergic neuronal diversity arises (Mayer et al., 2018; Mi et al., 2018) and to obtain an increasingly clear picture of how hypothalamic development is orchestrated (Kim et al., 2020; Romanov et al., 2020; Zhou et al., 2020). In spite of this progress, the molecular programs regulating the production of basal forebrain projection neurons in other regions remains poorly understood.

The septum is a ventral forebrain structure, mainly composed of a diverse array of GABAergic projection neurons that regulate a range of innate behaviors governing emotion and affect (Sheehan et al., 2004). The mature septum is segregated into a complex formed by the medial septum and the diagonal band of Broca (MS/DBB, hereafter referred to as ‘MS’ for simplicity) and lateral septum (LS) nuclei (in turn subdivided in dorsal, intermediate and ventral nuclei [LSd, LSi and LSv, respectively]), which have distinctive efferent and afferent connectivity with a variety of other areas of the brain (Sheehan *et al*., 2004). Most septal neurons are generated from neural progenitors located in two portions of the embryonic brain: the septum proper, which develops between the lateral ganglionic eminence (LGE) and the most anterior part of the cortex, and the septal eminence or ventral septum (Flames et al., 2007; Wei et al., 2012), a small and transient proliferative region adjacent to the medial ganglionic eminence (MGE) and rostral with respect to the embryonic preoptic area (PoA). The expression of Zic family transcription factors distinguishes septal progenitor zones from adjacent regions (Inoue et al., 2007). The developing septum can be further divided into pallial-like, LGE-like and MGE/PoA-like areas demarcated by the enriched expression of transcription factors such as *Tbr1*, *Gsh2* and *Nkx2.1*, respectively, which give rise to specific subpopulations of neurons (Flames *et al*., 2007; Puelles *et al*., 2000; Wei *et al*., 2012). Temporal production of neurons in the septum follows a general inside-out pattern: MS neurons are born earlier in development, while later-born cells occupy progressively more lateral positions in the LS (Wei *et al*., 2012). Despite some recent progress in understanding the spatial and temporal origins of diverse neuronal subgroups in the septum (Magno et al., 2017; Wei *et al*., 2012), little is known about the molecular programs guiding temporal competence states of septal progenitors and how they lead to the specification of different types of neurons. In this study, we address this by performing scRNA-Seq on septa at different stages in development to infer the developmental trajectories connecting progenitors to the neurons derived from them.

We generate a comprehensive dataset and interrogate it to gain insight into the extent neuronal diversity in the mature septum and how it is generated during development. We focus on two stages at the peak of MS and LS neurogenesis and find that the transcriptomic profile of neural progenitors and newborn neurons changes greatly between early and late neurogenic periods. Through genetic fate-mapping experiments, we resolve the contribution of the rostral and septal eminence to mature neuronal diversity. Finally, we describe the temporal pattern for the generation of diverse neuron types derived from the septal eminence defined by their morphological features and allocation within the septum. Distinctive subsets of septal neurons are important for regulating specific aspects of emotional and affective behavioral states. This study provides a comprehensive molecular framework for identifying the candidate gene networks involved in determining spatial and temporally defined septal neuron fates in a structure that is important for regulating emotion and affect.

## RESULTS

### Early emergence of distinct neuronal lineages in the developing septum

We selected six ages spanning the entire development of the septum, from the embryonic period to its maturity. We chose developmental stages that would cover the peak period of neurogenesis for both the medial (embryonic day [E]11) and lateral septum (E14) (Wei *et al*., 2012), as well as the subsequent processes of neuronal maturation, such as migration, axonal outgrowth and synaptogenesis (E17, postnatal days [P]3 and 10), along with P30 as a stage representative of a mature septum (**Figure 1A**, top). For each developmental stage, we manually dissected the septum (for E11, E14 and E17, the medial ganglionic eminences / preoptic areas, as potential sources of septal neurons (Wei *et al*., 2012), were collected as well), generated a single-cell suspension and subjected it to single-cell encapsulation followed by RNA sequencing, using a custom inDrops microfluidic system (Klein et al., 2015; Zilionis et al., 2017). After filtering the data to select for high-quality sequencing results (see **Methods**), we obtained a total of 72243 cells across our six developmental stages. Plotting the data using UMAP, a method that yields two-dimensional graphical representations where the position of individual cells conveys information about their relationships, we identified clusters corresponding to radial glia (RG), intermediate progenitors (IP) and neurons (N) based upon marker expression (**Figure 1B**). These identities were largely consistent with t-distributed stochastic neighbor embedding (t-SNE) of the same data (**Figure S1A**), where cluster identities were assigned based on an extensive complement of putative marker genes (**Figure S1B**). As expected from the experimental design, our dataset contains a number of distinct cell types, including different progenitor classes (radial glia and intermediate progenitors) and neurons at progressive stages of maturation (newborn, migrating, wiring and mature), as well as a number of other cells which were not the focus of this study (glia, ependymal cells, vasculature, etc.) (**Figure S1A,B**). However, in order to resolve potential relationships among individual cells and thus understand the molecular changes happening along defined lineages we needed a tool that allowed better visualization of inferred trajectories. We decided to use SPRING, a tool that generates a k-nearest-neighbor layout where each cell is represented as a node extending edges to the ‘k’ other nodes within the dataset with most similar gene expression profiles (Weinreb et al., 2017). This resulted in a graph containing all cells we sequenced, which were aligned according to their developmental stage despite the fact that SPRING was agnostic to the origin of each cell (**Figure 1C**). To keep the focus on septal neuron specification, we removed cells identified as non-neuronal (**Figure S1A,B**) and produced a SPRING plot containing 53011 cells that includes neurons at different maturation stages as well as the progenitors that gave rise to them (**Figure 1D**). We then produced SPRING plots of the E11 and E14 timepoints, corresponding respectively to the peak medial and lateral septal neurogenic periods, to identify the molecular pathways active during the specification of MS and LS neurons (**Figure 1E,F**). The genes *Nes* (nestin), *Ascl1* (achaete-scute family bHLH transcription factor 1), and *Dcx* (doublecortin), were used as cell type markers for radial glia, intermediate progenitors, and newborn neurons, respectively (Gleeson et al., 1999; Lendahl et al., 1990; Yun et al., 2002) (**Figure 1E,F**). The SPRING visualizations for E11 and E14 had a consistent organization, where more stem-like cells (i.e. radial glia) were located at one end of the plot, and more differentiated ones (i.e. neurons) at the opposite, with intermediate progenitors interspersed between them, reminiscent of the known lineage relationships among these three cell types (**Figure 1B,C**). Sectors of the SPRING plots comprised of newborn neurons were sharply divided into distinct protruding clusters, representing cells with common molecular identities. We assigned prospective identities to each of those clusters based upon their marker gene expression, confirming them based on the patterns of expression of each marker gene using the Allen Developing Mouse Brain Atlas (**Figures 1G,H; S1C, S2**). MGE/PoA-derived neurons were identified based on their expression of *Lhx6* (LIM homeobox 6) (Alifragis et al., 2004), while newborn cholinergic neurons expressed *Gbx2* (gastrulation brain homeobox 2) (Asbreuk et al., 2002; Chen et al., 2010); we also identified *Dner* (delta/notch like EGF repeat containing) as a general marker for septal neurons (**Figure 1G,H**). We identified additional distinct newborn neuronal populations, including presumptive pallium-derived neurons expressing *Tbr1* (T-box brain transcription factor 1) (Bulfone et al., 1995), septum-derived Cajal-Retzius cells (Bielle et al., 2005) expressing *Trp73* (transformation related protein 73) (Causeret et al., 2021; Meyer et al., 2004), and presumptive LGE-derived neurons expressing *Isl1* (ISL LIM homeobox 1) (Stenman et al., 2003; Toresson et al., 2000) (**Figure S2A,B**). The expression of these genes was restricted to the mantle zones of the MGE and the developing septum, further confirming that they are markers of postmitotic neurons, rather than progenitors (**Figures 1G,H; S1C**). While we recovered a relatively low number of cells from P30 samples (4145 cells; **Figure S3A**), and only a fraction (23 %) of these could be identified as MS or LS neurons, analysis of those groups of cells allowed us to identify several previously unreported potential markers for neurons located in either the LS or MS (**Figure S3B**), or common to both nuclei (**Figure S3C**). We identified *Prkcd* (protein kinase C delta) as a potential LS marker gene, and confirmed its restricted expression by crossing a Prkcd-Cre mouse line (Kalish et al., 2018) with a Cre-dependent reporter line (**Figure S3D**). Nearly all labeled cells were neurons confined to the lateral septum (**Figure S3E**), and thus could be assumed to be largely GABAergic (Zhao et al., 2013). As far as we are aware, this is the first report of a mouse line that grants wide genetic access to GABAergic neurons in the lateral septum. Together, these findings demonstrate that our scRNA-Seq dataset can be used to identify diverse molecular cell types and infer their developmental trajectories within the developing septum.

**Figure 1:**
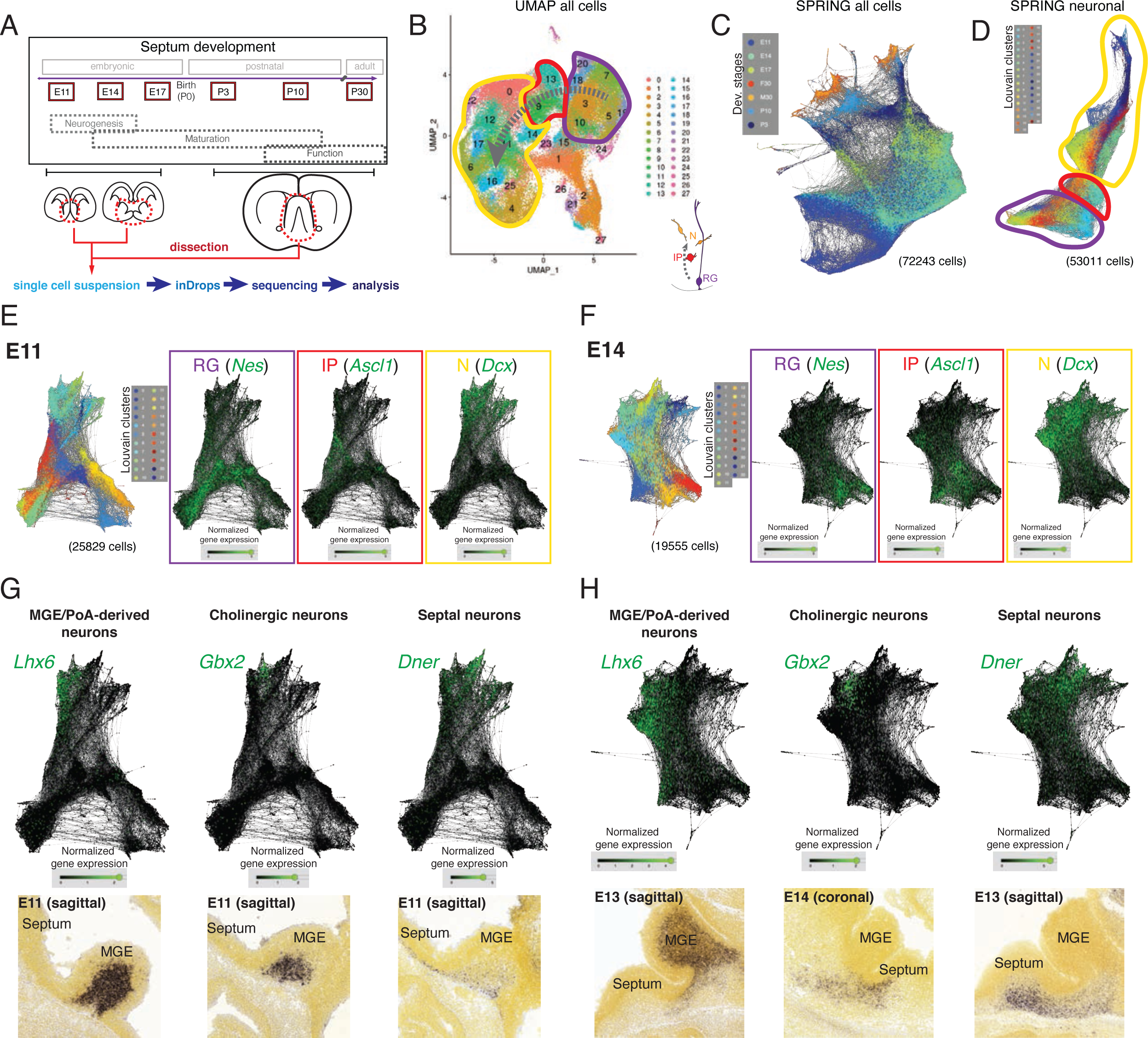
Single-cell RNA sequencing of the developing septum reveals early emergence of neuronal identities. **A)** Experimental approach: samples were collected at the indicated developmental stages and submitted to single-cell RNA-sequencing. **B)** UMAP plot of all cells, where the transition from radial glia (RG, purple) to neurons (N, yellow) via intermediate progenitors (IP, red) can be visualized (gray arrow). **C)** SPRING plot shows developmental stage-dependent organization of all sequenced cells. **D)** SPRING plot of cells belonging to neuronal trajectory. **E, F)** SPRING plots of all cells at embryonic stages E11 (E) and E14 (F) displays differentiation-dependent alignment of cells, illustrated by the relative enrichment of the genes *Nes*, *Ascl1*, and *Dcx* (cell type markers for RG, IP and N, respectively) in adjacent areas of the graphs. **G, H)** Analysis of each protrusion within the neuronal portion of E11 (G) and E14 (H) SPRING plots shows enrichment in marker genes for MGE/PoA-derived (*Lhx6*), cholinergic (*Gbx2*) and septal (*Dner*) newborn neurons, confirmed by *in situ* hybridization (bottom panels). Image credit (bottom panels): Allen Institute - Allen Developing Mouse Brain Atlas.

### Temporal transcriptional programs during MS and LS neurogenesis

Neurons destined for the medial septum are mainly generated during early neurogenesis, while those that will occupy the LS are born at later stages (Wei *et al*., 2012). We reasoned that since neuronal subtypes can already be distinguished at embryonic stages, different molecular programs guiding the generation of MS *versus* LS neurons from neural progenitor cells would be detectable as well. We compared the gene expression profiles of each of the three main cell types present along the neurogenic sequence (radial glia, intermediate progenitors, and newborn neurons) across the three embryonic stages, and found numerous genes that were differentially expressed (**Figure 2A**). Several classes of genes showed different levels of expression across developmental stages, including transcription factors, cell adhesion molecules, and intercellular and intracellular signaling molecules (**Figure S4A-D**). We focused on a select subset of genes composed of cell type markers and candidate temporal competence factors (**Figure 2B**). Since neurogenesis is largely completed by E17 and signatures of gliogenesis could already be detected, we decided to focus our subsequent analyses on the neurogenic stages E11 and E14. To validate our *in silico* differential gene expression analysis, we performed single-molecule *in situ* hybridization (RNAscope) for selected mRNA transcripts and quantified the signals during early (E12) and late (E14) septal neurogenesis (**Figure 2C-H**). We selected genes that were differentially expressed either at E11 or E14 and compared their expression in the septum proper (rostral) and the septal eminence (caudal) within the relevant zone demarcated by the expression of cell type marker genes (*Nes* for RG, *Ascl1* for IP, and *Dcx* for NN). Transcripts for genes of interest were normalized to the relevant marker gene. In RG, *Hmga2* was significantly enriched in E12 septum (**Figure 2C,F**), while *Hes5* was enriched at E14 (**Figure S4E**). In IP, both *Ccnd2* (**Figure 2D,G**) and *Ccnd1* (**Figure S4F**), were significantly enriched at E14. The gene *Prdm16*, a known marker of RG in other areas of the telencephalon (Baizabal et al., 2018; Shimada et al., 2017), was clearly upregulated in late-born neurons (**Figure 2E,H**); the same was true for other markers generally expressed at later neurogenic stages, such as *Nfia* (Clark et al., 2019) (**Figure S4G**). Together, our data provides a framework for characterizing dynamic gene expression as progenitors transition from generating MS neurons to LS neurons.

**Figure 2:**
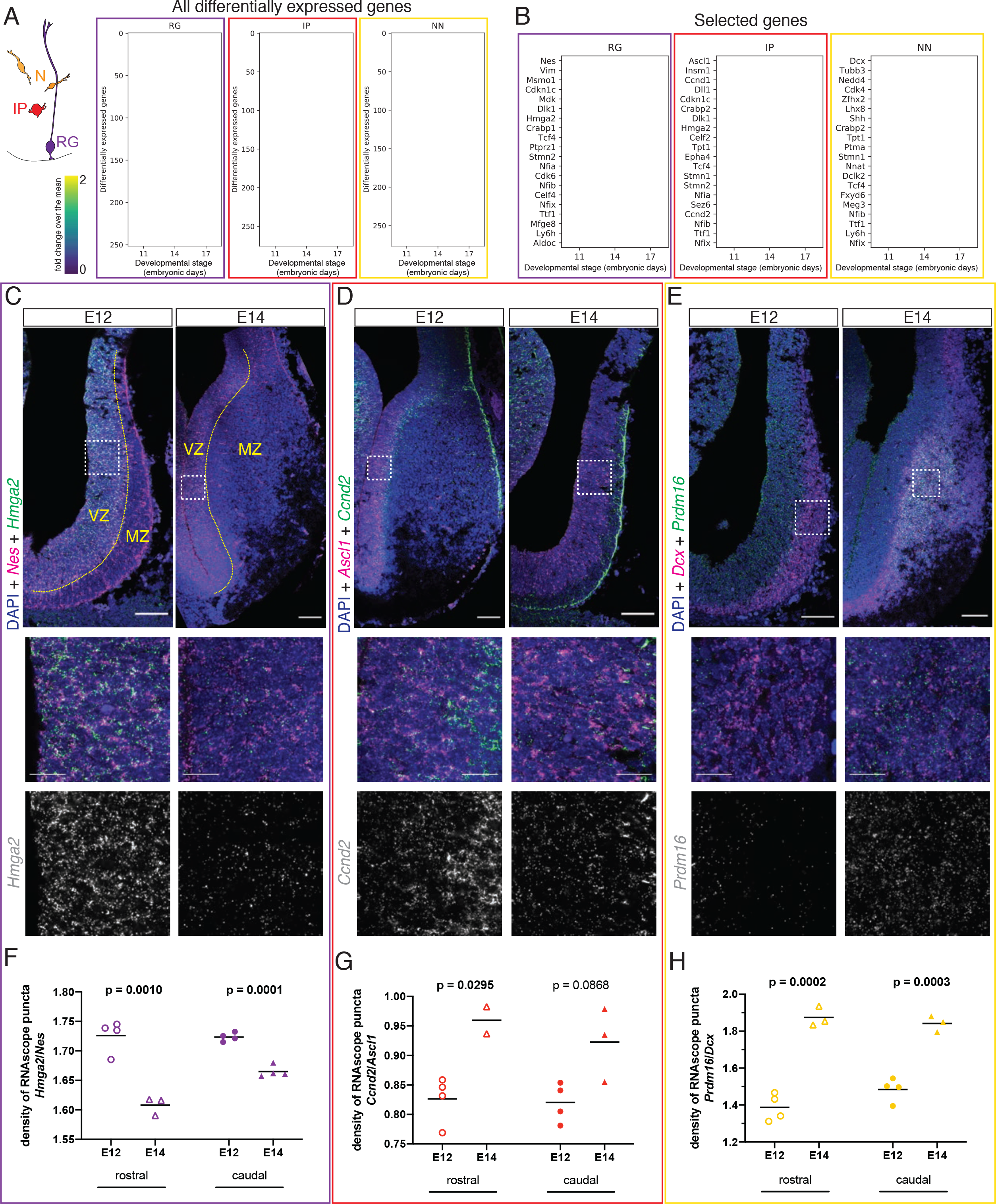
Different transcriptional programs are active during early and late septal neurogenesis. **A)** Heatmaps illustrating all genes differentially enriched in the scRNA- Seq dataset across embryonic stages for the three indicated cell types: radial glia (RG), intermediate progenitors (IP) and newborn neurons (NN). **B)** Heatmap showing the differential enrichment of a subset of selected genes, displayed as in A) and using the same color scale. **C-E)** Coronal brain sections showing the rostral septum at embryonic days 12 and 14, subjected to fluorescent single-molecule *in situ* hybridization for the genes *Nes* + *Hmga2* (C), *Ascl1* + *Ccnd2* (D) and *Dcx* + *Prdm16* (E). Cell type marker mRNA puncta (*Nes*, *Ascl1* and *Dcx*) are displayed in magenta, mRNA puncta for differentially enriched genes (*Hmga2*, *Ccnd2* and *Prdm16*) in green, and DAPI counterstaining in blue. Yellow dashed lines in C) mark the limit between ventricular zone (VZ) and mantle zone (MZ), as indicated. Dashed boxes indicate the location of the magnified 100x100-µm fields shown below, both as merged images (middle panels) and single-channel for the corresponding differentially enriched genes (bottom panels). Scale bars: 100 µm (top panels), 25 µm (middle panels). **F-H)** Quantification of density of mRNA puncta of differentially enriched genes, normalized to the density of cell type marker mRNA; measurements were obtained from the rostral (empty symbols) and caudal (full symbols) portions of the septum at E12 (circles) and E14 (triangles). All data points are represented; black bars represent the mean. Unpaired t-tests were performed; p-values are indicated above the corresponding compared sets of data: those highlighted in bold indicate statistically significant differences (p<0.05).

### Neurons in both the MS and LS are derived from the septal eminence

Previous work has shown that a portion of cells in the septum, most notably MS cholinergic neurons, are derived from progenitors expressing the transcription factor *Nkx2.1* (Magno *et al*., 2017; Wei *et al*., 2012; Xu et al., 2008). Our embryonic samples contained cells from both the developing septum and the MGE/PoA, which could be identified in our SPRING plots (**Figure 1G,H**). E14 SPRING plots of *Nkx2.1* and *Zic4*, a general septal marker (Rubin et al., 2010) revealed a region of overlapping expression spanning across zones containing progenitors and newborn neurons (**Figure 3A**). We confirmed that embryonic progenitors located in the VZ/SVZ proliferative areas of the caudal, but not the rostral, septum were positive for NKX2.1 (Magno *et al*., 2017) (**Figure 3B**), and found a stream of ZIC-positive cells with a history of *Nkx2.1* expression migrating rostrally from their putative site of origin in the septal eminence (**Figure 3C**). To better understand the contribution of rostral and caudal septal progenitors to septal neuron diversity, we used three different genetic mouse models to fate-map cells derived from *Zic4*-expressing (all septal proliferative areas), *Nkx2.1*- expressing (MGE and septal eminence) or *Zic4* and *Nkx2.1* co-expressing (septal eminence) progenitors (**Figure 3D-F**). The vast majority of neurons (71-97%) in the three main subdivisions of the LS (dorsal, intermediate and ventral nuclei), and approximately 50% of MS neurons, had a history of *Zic4* expression throughout the rostro-caudal axis (**Figure 3G**). About 10-30% of ZIC+ cells in both the LS and the MS (in a decreasing proportion along the rostro-caudal axis) had a history of *Nkx2.1* expression (**Figure 3E**); to confirm that these were derived from septal rather than MGE/PoA progenitors, we used an intersectional genetic approach, combining an Nkx2.1-Flp knock-in driver line with a Zic4-Cre driver line and the FLTG reporter line (Plummer et al., 2015). Cells with a history of expression of both genes would result in labeling with GFP, while expression of only *Nkx2.1* would label cells with tdTomato (Plummer *et al*., 2015). The vast majority of labeled cells in the septum, with the exception of an abundant MS astrocyte population, were positive for GFP, confirming their origin in the septal eminence (**Figure 3F,I**). The distinct neuronal output from the septal eminence and its expression of *Nkx2.1* suggests that there may be fundamental differences in progenitor composition and proliferative behaviors between the rostral and caudal proliferative zones in the embryonic septum (Magno *et al*., 2017). To address these potential differences, we performed immunostaining for the mitotic marker phosphorylated histone 3 in E13 samples in order to compare the abundance and location of cycling progenitors in the developing septum and the septal eminence (**Figure S5A**). This experiment revealed a much lower proportion of dividing cells located in the putative subventricular zone of the septal eminence, when compared to the rostral portion of the septum (**Figure S5A,B**). This could reflect underlying cell biological differences between these two regions, whereby fate-restricted *Ascl1*- expressing progenitors in the septal eminence would preferentially undergo direct neurogenic divisions at the ventricular surface rather than delaminating and entering the subventricular zone transit amplifying intermediate progenitors (Petros et al., 2015; Turrero Garcia and Harwell, 2017). This was supported by a roughly two-fold increase in the proportion of ASCL1+ mitotic cells at the ventricular surface of the NKX2.1+ portion of the septal anlage (**Figure S5C,D**). Together, these data suggest that the septal eminence is composed of progenitors with distinctive molecular profiles, proliferative capacity and fate potential compared to the rostral embryonic septum.

**Figure 3:**
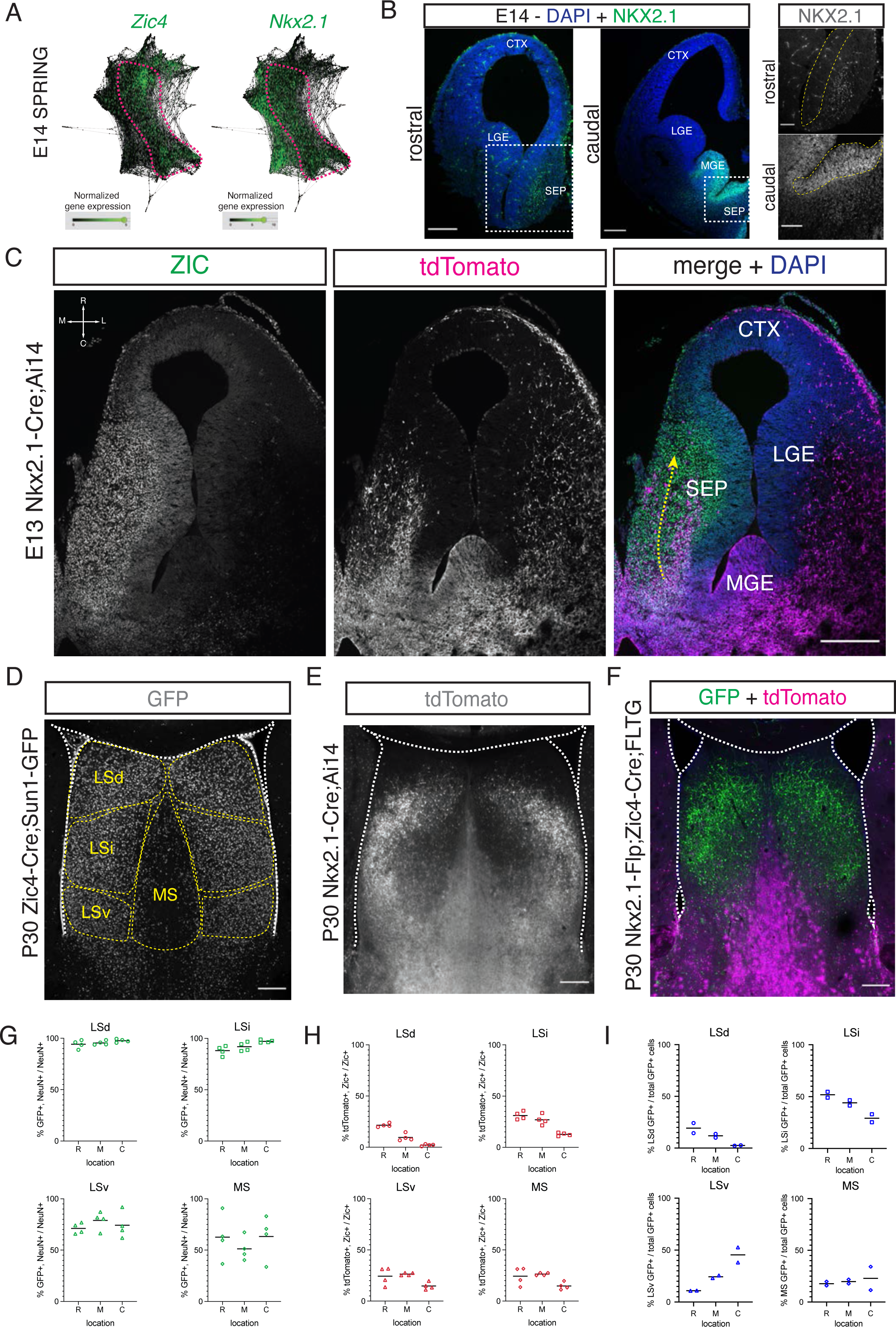
Contribution of the septal eminence to neuronal composition of the mature septum. **A)** SPRING plots at E14 show that a set of cells at this stage (dashed magenta line) express both *Zic4* (left) and *Nkx2.1* (right). **B)** Immunofluorescence staining for NKX2.1 (green; counterstained with DAPI, blue) at rostral and caudal locations of the septum on coronal sections of an E14 brain. Panels on the right display magnified insets (marked with dashed white squares on the corresponding overview images); the main proliferative area (ventricular zone) is highlighted by a yellow dashed line, showing NKX2.1-positive cells in the caudal, but not rostral, septal anlage. Scale bars: 250 µm (overview images); 100 µm (closeup images). **C)** Horizontal section (compass on the top left of left panel indicates rostro-caudal and medio-lateral axes) of the right hemisphere of an E13 Nkx2.1-Cre;Ai14 mouse brain. Immunofluorescence staining for ZIC proteins (left panel; green in merge) and the tdTomato fluorescent reporter (middle panel; magenta in merge), shown both as single channels and merged, (right panel, with DAPI counterstaining in blue) show a subset of *Nkx2.1*-expressing septal progenitors in the septal eminence, as well as a caudal-to-rostral stream of migrating ZIC-positive neurons with a developmental history of *Nkx2.1* expression (yellow dashed arrow). Scale bar, 250 µm. **D)** Coronal section of the septum of a P30 Zic4-Cre;Sun1-GFP mouse; immunofluorescence staining for the reporter GFP shows the location of *Zic4*-lineage cells. Yellow dashed lines indicate the location of MS and LS nuclei. **E)** Coronal section of the septum of a P30 Nkx2.1-Cre;Ai14 mouse; immunofluorescence staining for the reporter tdTomato shows the location and morphology of *Nkx2.1*-lineage cells. **F)** Coronal section of the septum of a P30 Nkx2.1- Flp;Zic4-Cre;FLTG mouse; immunofluorescence staining for the reporters GFP (green) and tdTomato (magenta) shows the location and morphology cells within the *Nkx2.1*- lineage with (GFP+) or without (tdTomato+) a developmental history of *Zic4* expression. Scale bars (D-F), 250 µm. **G-I)** Quantification of the proportion of neurons within the *Zic4*-lineage within the total NeuN+ neuronal population (G), the proportion of *Nkx2.1*- lineage cells within the ZIC+ population (H), and the proportion of the entire intersectional (GFP+) population within each septal nucleus (I), obtained for each of the mouse lines above the corresponding set of graphs, across the dorsal, intermediate and ventral nuclei of the lateral septum (LSd [circles], LSi [squares] and LSv [triangles], respectively) and the medial septum (MS [diamonds]) at rostral (R), medial (M) and caudal (C) locations along the antero-posterior axis. Abbreviations: CTX: cortex; LGE: lateral ganglionic eminence; MGE: medial ganglionic eminence; SEP: septum.

### Morphology and distribution of temporal cohorts of neurons in the *Nkx2.1*-lineage

Septal neurons are born in a defined inside-out temporal sequence, where the MS is generated early in neurogenesis and later-born neurons are allocated to progressively more lateral positions (Wei *et al*., 2012). To better understand the patterning of temporal fates of neurons derived from the septal eminence, we devised an intersectional approach based on the *Ascl1* expression by temporally fate restricted neurogenic progenitors within the *Nkx2.1*-lineage (Kelly et al., 2019) (**Figure 4A**). We combined the Nkx2.1-Flp driver mouse line with a tamoxifen (TMX)-inducible Ascl1-CreER line and the intersectional Ai65 reporter line, which expresses the fluorescent protein tdTomato in a Cre- and Flp-dependent manner (**Figure 4B**). Administration of tamoxifen causes the expression of tdTomato in cells with a developmental history of *Nkx2.1* expression that are undergoing a peak of *Ascl1* expression, leading to a neurogenic division. We administered tamoxifen to timed-pregnant dams at four embryonic timepoints spanning the neurogenic period (E10, E12, E14 and E16), and collected the brains of their progeny at P30, when development of the septum is complete (**Figure 4C**). Temporally-defined cohorts of tdTomato-expressing cells were distributed primarily within the lateral septum (**Figure 4D**). Cells labeled at different stages were not distributed in a strictly inside-out pattern, but rather in a roughly medial-to-dorsal-to-ventral one. Considering the temporal dynamics and viewing each temporal cohort as a still picture within a sequence, this pattern can be compared to water in a fountain or a fireworks display, whereby cells are initially located in the center of an upwards stream, “moving” into more dorsal positions until they reach the apex and then “falling on the outside” to progressively more ventral positions (**Figure 4D,E, S6C**). We adopted and extended the classification morphological types of LS neurons described by Alonso and Frotscher in 1989 (Alonso and Frotscher, 1989) to determine the extent of morphological diversity within each neuronal temporal cohort. We found the same morphological neuronal types regardless of their location within the different nuclei within the LS, We therefore designated labeled cells as type I, II and III, irrespective of their allocation to LSd, LSi or LSv, as follows (see **Methods** for further details): type I neurons, with relatively few thick dendrites forming a spherical contour; type II neurons with thinner and branched dendrites; we propose to further subdivide the morphological types by adding a neuronal type III, with thick, spine-dense dendrites forming a bipolar dendritic field (**Figure 4F**). With the exception of E16, where only a few mostly type II cells were present in the LSv, there were neurons of all three types in each temporal cohort labeled, with slight changes in their overall distribution (**Figure 4G**). Type I cells were most abundant in the LSd; type II neurons were prevalent across all LS areas and temporal cohorts, and the proportion of type III cells was highest in the E14 cohort, and practically confined to LSd and LSi (**Figure S6A**). Type II neurons could be further classified into three subgroups based upon the morphology of their dendrites; type IIa, with overall thicker dendrites; type IIb with a thick initial dendritic segment bifurcating into thinner processes, and type IIc, with thin and long dendrites (**Figure S6B**). This set of experiments demonstrates that while septum development generally follows the inside-out model previously described (Wei *et al*., 2012), certain subsets of neurons, such as those derived from *Nkx2.1*-expressing progenitors in the septal eminence, might follow different patterns. We expand upon the morphological classes proposed by Alonso and Frotscher, describing the newly defined type III neurons. The LSd/LSi location and bipolar shape of the dendritic field of type III neurons are likely reflective of unique functional/connectivity properties, which will require further investigation (**Figure S6C**).

**Figure 4:**
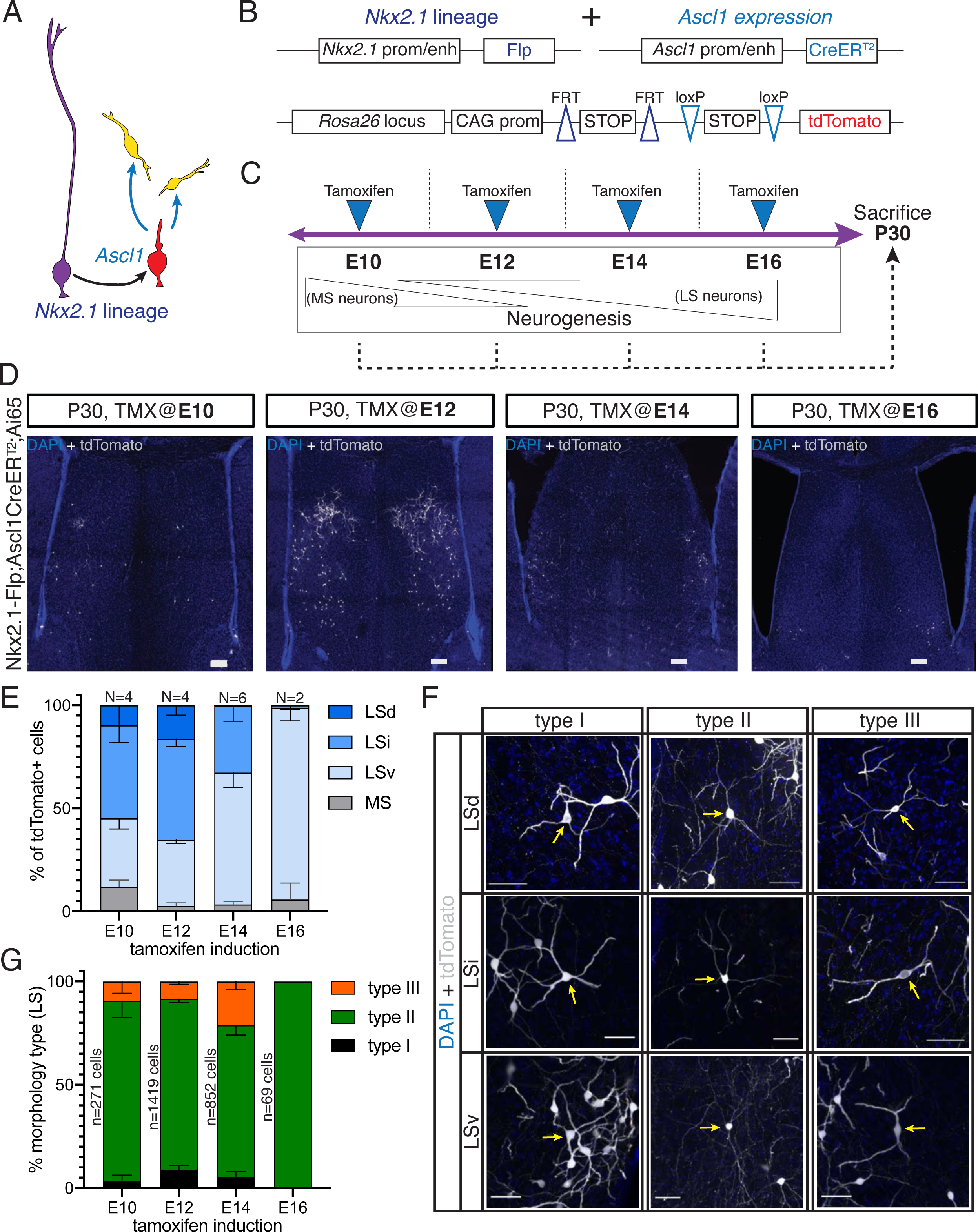
Generation of sequential temporal cohorts of neurons from the septal eminence. **A)** Schematic of neurogenesis in the septal eminence: RG (purple) within the *Nkx2.1* lineage give rise to *Ascl1*-expressing transit-amplifying progenitors (red), which in turn divide to generate neurons. **B)** Two driver lines, Nkx2.1-Flp and Ascl1- CreER^T2^, were crossed with an intersectional reporter line. The action of both Flp and Cre recombinases (i.e., within the *Nkx2.1* lineage and in the presence of tamoxifen) leads to the expression of the fluorescent reporter tdTomato. **C)** Experimental design: tamoxifen was administered to pregnant dams at E10, E12, E14 or E16, covering the entire septal neurogenic period. Resulting litters were sacrificed and analyzed at P30. **D)** Representative images of coronal sections through the septum of P30 mice in which recombination was induced at the indicated stages. Cells derived from terminal progenitor divisions are labeled with tdTomato (gray; counterstained with DAPI, blue). Scale bars, 100 µm. **E)** Quantification of the location of tdTomato+ cells in each septal nucleus, as a percentage of total labeled cells (mean ± s.d.) within each temporal cohort. The number of biological replicates is indicated above each bar. **F)** Representative images of neurons belonging to the three morphological subtypes within each LS nucleus, labeled with tdTomato (gray; counterstained with DAPI, blue). Arrows indicate the cell bodies of corresponding neuronal types. Scale bars, 50 µm. **G)** Quantification of morphological neuron types, as a percentage of the total number of classified cells (mean ± s.d.) within each temporal cohort. The total number of classified cells is indicated on the left side of the corresponding column; the number of biological replicates is the same as in E except for the E14 timepoint (N=7).

## DISCUSSION

The full diversity of neurons contained within the septum and the dynamic transcriptional programs controlling their generation from embryonic progenitors are still largely unexplored. Foundational work from the Yang lab explored a number of neuronal subpopulations in the mature septum, demonstrating that they were derived from multiple progenitor regions located within the septum itself, the MGE/PoA, and the pallium or pallial/subpallial boundary (Wei *et al*., 2012). These findings were extended with the development of a septal-specific Zic4-Cre driver mouse line (Rubin *et al*., 2010), which allowed for a more refined molecular classification of septal progenitor zones. The rostral and caudal portions of the developing septum can be divided into LGE-like and the MGE-like progenitor zones, respectively. While the rostral septum and the LGE serve as the anatomical templates for their mature derivatives (the septum proper and the striatum), the MGE and the septal eminence are largely transient structures with merely vestigial counterparts in the mature brain (Delgado and Lim, 2017; Delgado et al., 2020; Merkle et al., 2014). The main role of the MGE and the septal eminence during embryonic development is to generate cells destined to occupy other areas of the brain (cortical interneurons in the MGE, multiple septal neurons and basal forebrain cholinergic neurons in the septal eminence). Both of these proliferative areas express *Nkx2.1*, a gene that is fundamental to maintain their correct regional identity and neuronal output.

Our work helps to clarify the contribution of the septal eminence to the diversity of neurons in the medial and lateral nuclei of the mature septum. Consistent with previous reports, we found that the vast majority of neurons throughout the lateral septum, and about half of those in the medial septum, belong to the *Zic4* lineage (**Figure 3D,G**). Roughly a quarter of those neurons have a developmental history of *Nkx2.1* expression (**Figure 3E,H**), and could thus originate from either the septal eminence or the MGE/PoA, as has been proposed before (Wei *et al*., 2012). To resolve this, we used an intersectional genetic approach to label cells expressing both *Nkx2.1* and *Zic4*, confirming that the vast majority of *Nkx2.1*-lineage neurons in the MS and LS are derived from the septal eminence rather than from the MGE/PoA, since these regions do not express *Zic4* (Magno *et al*., 2017). More work will be necessary to understand how the interplay of these and other genes with regionally restricted expression patterns, such as fibroblast growth factors (Hoch et al., 2015), impact the neuronal output of either septal progenitor zone. We hypothesize that further subdivisions within the developing septum (Flames *et al*., 2007) are responsible for the generation of specific neuronal subtypes, as in the MGE (Hu et al., 2017).

Our data suggest that the proliferative areas of the septum proper and the septal eminence have diverging molecular identities, which likely lead to differences between the cell biology of the progenitors contained within them, in terms of both cell type diversity and proliferative behavior. We observed that progenitors in the septal eminence divide in more apical locations than their rostral counterparts (**Figure S5A,B**), and that a higher proportion of ventricular divisions are ASCL1-positive (**Figure S5C,D**). This suggests that *Nkx2.1*-expressing septal eminence progenitors may undergo transit-amplifying cell divisions as short neural precursors rather than delaminated intermediate progenitors (Petros *et al*., 2015; Turrero Garcia and Harwell, 2017). Future work comparing the specific cell biological features of septal progenitors located within defined subdomains should help to elucidate how cell fate specification occurs at the cellular and/or clonal level (Zhou *et al*., 2020).

Temporal changes in the transcriptional profiles of ventral telencephalic progenitors guide their competence to generate different types of neurons as neurogenesis progresses (Turrero Garcia and Harwell, 2017). We hypothesized that similar changes would underlie the inside-out patterning of the septum. To test this, we used our scRNA-Seq dataset to compare gene expression levels across developmental stages and cell types (**Figure 2**). Several of the genes we validated recapitulate temporal patterns of expression that have been described in other parts of the developing central nervous system. For example, *Hmga2* is highly expressed in early stages of neurogenesis across all septal cell types we analyzed (**Figures 2C,F, S4B**); this is similar to cortical progenitors, where it controls early developmental programs (Shu et al., 2019). Likewise, we observed an increase in the level of expression of *Ccnd2* (**Figure 2D,G**) in intermediate progenitors at later developmental stages, which suggests that its role in cell cycle regulation and its interplay with *Ccnd1* (**Figure S4F**) are crucial for ensuring the correct neuronal outputs of late septal progenitors, as it is in the MGE (Glickstein et al., 2007; Glickstein et al., 2009). We also found a general upregulation in Nfi factors (**Figures 2B, S4G**) during late stages of septal neurogenesis, reminiscent of their role in late fate specification in the retina (Clark *et al*., 2019). Further research will be necessary to determine whether these or other temporally enriched factors promote the specification of specific septal neuron fates. While location and time of progenitor divisions appear to determine the general identity of septal neurons (**Figures 1G,H; S2; 3**), it is likely that the confluence of extrinsic factors and cell-cell interactions further refines their fate specification (Fishell and Kepecs, 2020).

Another aspect we considered when analyzing patterns of gene expression was the progression of neural progenitors from radial glia to neurons within each developmental stage. We noticed that *Prdm16* is highly expressed in both RG and newborn neurons, especially late-born (**Figure 2E,H**). In other parts of the developing brain, *Prdm16* is expressed exclusively in radial glia, and quickly shut down as cells become intermediate progenitors (Baizabal *et al*., 2018; Turrero Garcia et al., 2020). Its upregulation to even higher levels in septal neurons could reflect a novel role for this gene in the control of further aspects of neuronal differentiation, especially in the LS. Other transcription factors such as *Sp8*, *Pax6*, *Sox6*, or *Nkx2.1*, which are usually downregulated by postmitotic neurons in other parts of the brain, maintain high levels of expression in septal neurons into adulthood (Wei *et al*., 2012). While the role of several of these genes has been explored in postmitotic MGE-derived neurons (Azim et al., 2009; Batista-Brito et al., 2009; Nobrega-Pereira et al., 2008), their continued expression in septal neurons has not been addressed so far. This phenomenon could be an intrinsic part of the process of septal neuron fate specification and/or maintenance of subtype specific neuronal identity and circuitry(Sheehan *et al*., 2004).

Earlier studies have analyzed the birthdates of mature septal neurons expressing specific markers, and thus assumed to share a common origin (Wei *et al*., 2012). Here, we devised a genetic strategy to label isochronic temporal cohorts of septal eminence-derived neurons (**Figure 4A-C**). We found that cells within the *Nkx2.1*-lineage are largely located in the LS, and generated in a dorsal-to-ventral, rather than medial-to-lateral, gradient (**Figure 4C,D**). Our labeling strategy allowed us to study the morphology of these neurons (**Figure 4E**) and the distribution of defined morphological subtypes (**Figures 4F, S6A**). We based our analyses in the foundational work of Alonso and Frotscher (Alonso and Frotscher, 1989), with three additions: 1) since the morphological classes within the *Nkx2.1*-lineage were not clearly segregated across LSd and LSi (**Figure S6A**), we decided to unify the nomenclature, basing it in the type I and type II categories described for the LSd and extending it to the LSv as well; 2) we consistently observed neurons with thick dendrites and bipolar shape, which we designated as type III septal neurons (**Figure 4E**); and 3) given the morphological variability within type II cells, we propose to further classify them into subtypes IIa, IIb and IIc (**Figure S6B**). It is possible that neurons derived from the septal eminence with additional morphologies, corresponding to other types described by Alonso and Frotscher (Alonso and Frotscher, 1989). However, we examined >2600 cells labeled throughout the entire septal neurogenic period (**Figure 4G**), which suggests that, unless there are cells that our analyses might have missed (such as neurons born from direct neurogenic divisions, bypassing the *Ascl1*-expressing stage), we have captured the diversity of LS *Nkx2.1*-lineage neurons. One important caveat of our current approach is that we have not been able to correlate morphological types to transcriptional profiles, beyond the fact that they would belong to a LS group (clusters 2, 5 and 7 in **Figure S3A**). This is due in great part to the relative scarcity of our mature septum dataset, and should be addressed by future experiments where a higher resolution can be achieved. It will be important to assess if and how these morphological types differ in terms of their molecular identity and connectivity patterns in order to better understand their function within septal circuits and consequently their role in behavioral regulation. It will be particularly interesting to study how the three morphological types we have observed might be correlated to the three classes of LS neurons that have been described based on their electrophysiological properties (Gallagher et al., 1995; Wang et al., 2019).

Single-cell sequencing techniques allow unprecedented interrogation of the molecular diversity of cell types present in any tissue. Here, we provide the first scRNA-Seq dataset of the developing and mature septum that we are aware of, in a format that will allow other investigators to use it as a springboard towards further discoveries. Since we included the MGE/PoA in our dissections, our data can be used to complement and extend previously published scRNA-Seq datasets addressing cell diversity in this area (Mayer *et al*., 2018; Mi *et al*., 2018). Our current analysis highlights the point in the molecular trajectory of septal progenitors when they acquire distinct states that are predictive of their cardinal neuronal subtype identity (**Figure 1G,H)**. Our data suggests the mechanism for determination of cardinal cell type identity is similar to what is observed in neighboring structures such at the MGE (Fishell and Kepecs, 2020).

The initial exploration of P30 samples within our dataset has yielded several previously undescribed markers of neuronal subpopulations in the adult septum (**Figure S3A-C**), one of which we validated with a transgenic mouse line that grants access to GABAergic neurons in the LS (**Figure S3D,E**). However, this study focuses largely on developmental stages; future research efforts should address mature septal neuronal diversity in a more systematic and comprehensive way. The septum is involved in numerous psychological and psychiatric conditions, including psychotic spectrum disorders, anxiety and depression (Sheehan *et al*., 2004), Despite this, very few detailed descriptions of the human septum (Andy and Stephan, 1968) or its development (Rakic and Yakovlev, 1968) have been published. Considering the high evolutionary conservation of septal nuclei across tetrapods in terms of both anatomy and function (Lanuza and Martínez-García, 2009), studies like ours are likely to shed light on common mechanisms of cell fate determination, and uncover species specific cell types and developmental programs. More detailed scRNA-Seq experiments across multiple species, including humans, will be necessary to fully understand how septal neuronal diversity is specified during development, and how it impacts brain function and behavior.

## MATERIALS AND METHODS

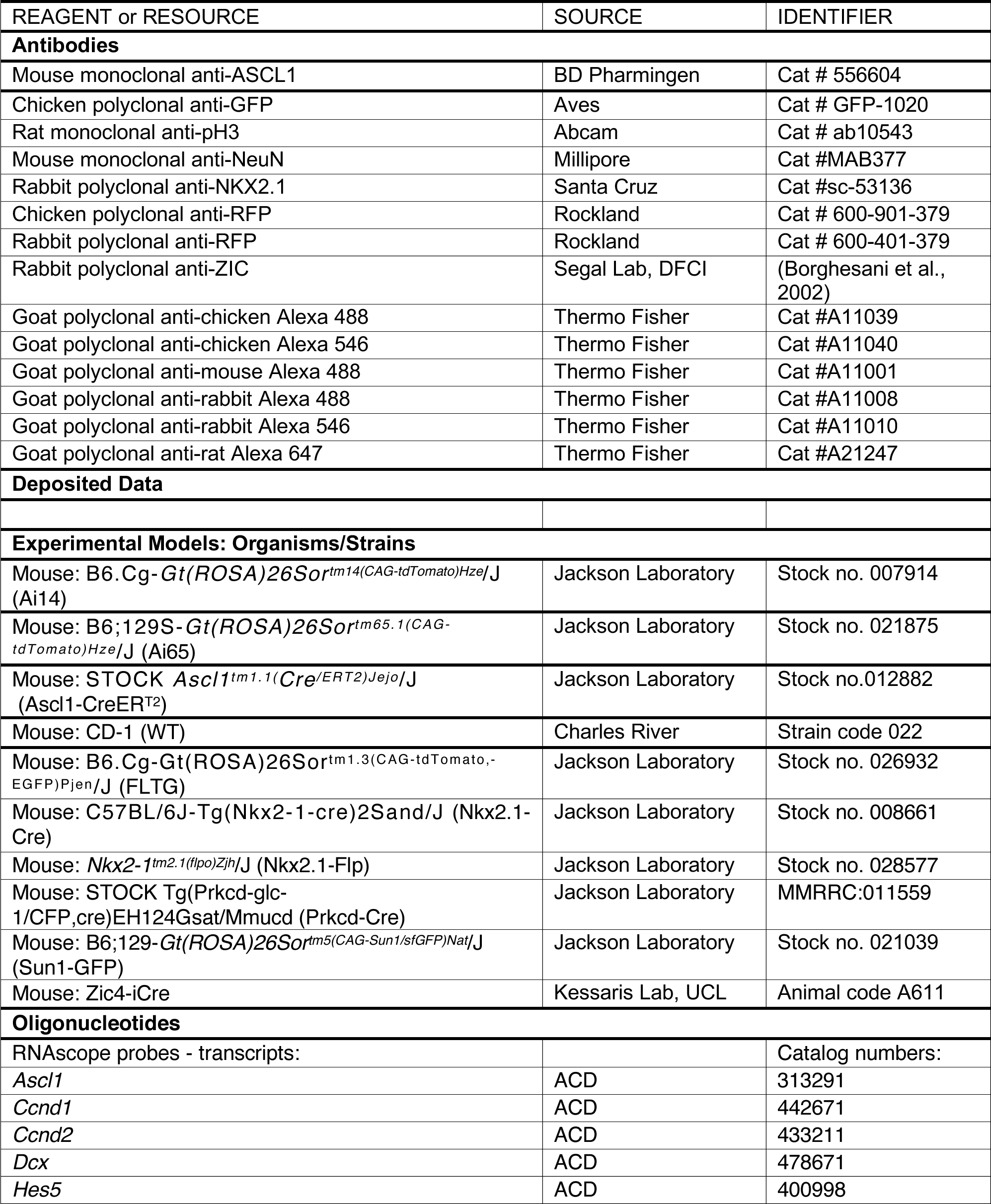
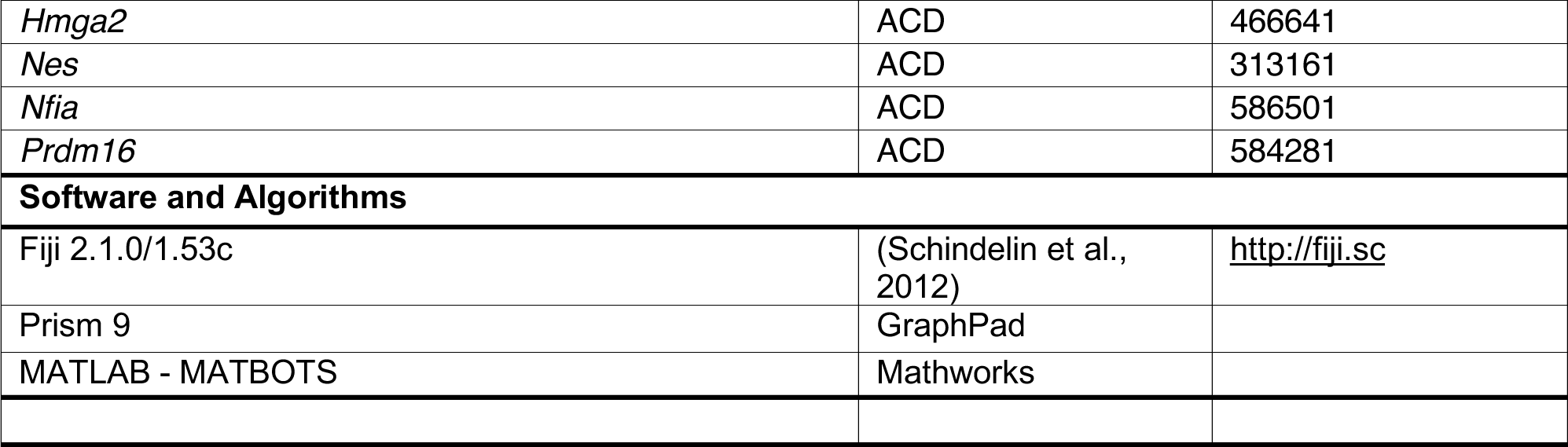
KEY RESOURCES TABLE.

### EXPERIMENTAL MODEL AND SUBJECT DETAILS

All animal procedures conducted in this study followed experimental protocols approved by the Institutional Animal Care and Use Committee of Harvard Medical School. Mouse lines are listed in the Key Resources table. Wild-type animals for single cell sequencing and validation experiments were purchased as timed-pregnant females or as entire litters at the following developmental time points: embryonic days (E)11, 14 and 17; and postnatal days (P)3, 10, and 30. For all other experiments, mouse housing and husbandry were performed in accordance with the standards of the Harvard Medical School Center of Comparative Medicine. Mice were group housed in a 12 h light/dark cycle, with access to food and water *ad libitum*. Samples were obtained at the ages indicated in the Figure Legends and throughout the text; for embryonic samples, the plug date was considered as E0. All results reported include animals of both sexes, balanced equally wherever possible; the sex of embryos and P3 animals was not determined. The number of animals used for each experiment (i.e. biological replicates) is indicated in the corresponding graphs where possible. The number of animals used for scRNA-Seq experiments was: E11 – 35 embryos from 3 litters; E14 – 33 embryos from 3 litters; E17 – 8 embryos from 2 litters; P3 – 4 pups from 2 litters; P10 – 2 pups (1 male, one female) from 1 litter; P30 – 2 males and 2 females from 1 litter.

## METHOD DETAILS

### Dissection, single cell suspension and droplet capture

For embryonic samples, the pregnant dam was sacrificed; embryos were removed from the uterus and maintained in Hibernate-E medium minus CaCl2 (HEMC), on ice, for the remainder of the procedure. Postnatal animals were deeply anesthetized and transcardially perfused with ice-cold PBS. Brains were extracted and transferred to dissection medium (HEMC + 0.1 mg/ml DNAse I). All septa (medial ganglionic eminences were also collected from embryonic samples) were manually dissected out and lightly minced with the dissection forceps, then transferred with a minimum amount of medium to an Eppendorf tube containing 1 ml of Accutase and 0.1 mg/ml of DNAse I. They were then incubated, rocking at 4°C, for 10-15 min. The tube was then centrifuged at 1000 rpm, at 4°C, for 1-2 min. After discarding the supernatant, the tissue was resuspended in 1 ml of dissection medium, and dissociated by gently pipetting up and down, first with a 1000 µl pipette tip (10-15 times), and subsequently with a 200 µl tip (10 times), both loaded to half to two-thirds capacity. The suspension was centrifuged at 1000 rpm, at 4°C, for 5 min. The supernatant was discarded, and cells were resuspended in 1 ml of HEMC and filtered through a 35-µm cell strainer and transferred to a clean low-adhesion Eppendorf tube. The resulting single-cell suspension was maintained on ice and subjected to single-cell droplet encapsulation with a custom microfluidic inDrops system at the Harvard Medical School Single Cell Core. Cell encapsulation and library preparation followed a previously described protocol (Klein *et al*., 2015; Zilionis *et al*., 2017), with modifications in the primer sequences as included in the Key Resources table.

### Single-cell RNA sequencing

Libraries of approximately 3,000 cells were collected from each animal. inDrops was performed as previously described (Hrvatin et al., 2018; Klein *et al*., 2015; Zilionis *et al*., 2017), generating indexed libraries that were then pooled and sequenced across 8 runs on the NextSeq 500 (Illumina) platform.

### inDrops sequencing data processing

Transcripts were processed according to a previously published pipeline (Hrvatin *et al*., 2018; Klein *et al*., 2015; Zilionis *et al*., 2017). Briefly, this pipeline was used to build a custom transcriptome from the Ensembl GRCm38 genome and **GRCm38.88** annotation with Bowtie 1.1.1 (after filtering the annotation gtf file (ftp://ftp.ensembl.org/pub/release-88/gtf/mus_musculus/Mus_musculus.GRCm38.88.gtf.gz filtered for feature_type = ‘gene’, gene_type = ‘protein_coding’ and gene_status = ‘KNOWN’). Read quality control and mapping against this transcriptome were then performed. Finally, unique molecular identifiers were used to reference sequence reads back to individual captured molecules, thus yielding values denoted as UMI counts. All steps of the pipeline were run with default parameters unless explicitly specified.

### Quality control for cell inclusion

Cells from each time point (E11, E14, E17, P3, P10, P30) were pre-processed separately. Any cells with fewer than 700 or more than 10000 transcript counts were excluded from the analysis. Any cells in which >50% of UMIs mapped to mitochondrial genes were excluded. The dataset was normalized (NormalizeData()), variable genes identified (FindVariableGenes(x.low.cutoff = 0.0125, x.high.cutoff = 3, y.cutoff = 0.5)). The data was scaled using variable genes and a negative binomial model with the percentage of mitochondrial genes and the number of UMIs per cell regressed (ScaleData(vars.to.regress = c(“percent.mito”, “nUMI”), genes.use =, model.use = “negbinom”)). PCA analysis, clustering, tSNE plotting and marker identification were performed using recommended parameters: RunPCA(pc.genes =, pcs.compute = 40, pcs.print = 1:30, maxit = 500, weight.by.var = FALSE); FindClusters(dims.use = 1:30, resolution = 1.5, print.output = 1, save.SNN = T,reduction.type = “pca”); RunTSNE(dims.use = 1:30, do.fast = T); FindAllMarkers(only.pos = F, min.pct = 0.1, thresh.use = 0.25). By inspection of the tSNE plots and marker genes, one to three clusters were identified at each timepoint as likely doublet clusters and those cells were excluded from further analysis. Our dataset after quality control contained 72,243 cells with more than 700 reads assigned to each cell.

### Dimensionality reduction and clustering

All 72,243 cells were combined into a single dataset and analyzed simultaneously. The R software package Seurat (Butler et al., 2018; Satija et al., 2015) was used to cluster cells. First, the data were log-normalized and scaled to 10,000 transcripts per cell. Variable genes were identified using the FindVariableGenes() function. The following parameters were used to set the minimum and maximum average expression and the minimum dispersion: x.low.cutoff = 0.0125, x.high.cutoff = 3, y.cutoff = 0.5. Next, the data was scaled using the ScaleData() function, and principle component analysis (PCA) was carried out. The FindClusters() function using the top 30 principal components (PCs) and a resolution of 1.5 was used to determine the initial 32 clusters.

## SPRING

SPRING plots were generated using the standard SPRING pipeline (Weinreb et al., 2018) with modifications described in (Weinreb et al., 2020). Briefly, UNI counts were total counts normalized (without log-normalization) and filtered for highly variable genes. Gene expression values were standardized to zero-mean and unit-variance, and a low-dimensional embedding was estimated with Principal Components Analysis (PCA). The final 2D layout was produced by applying the ForceAtlas2 graph layout algorithm to a k-nearest-neighbor graph over PCA coordinates with k=3.

### Bioinformatic analyses – cluster ID and differential gene expression

Clusters were assigned a cell type label by manual inspection of marker gene expression. We curated a list of known markers from the literature and constructed as cluster-by-marker heatmap as follows: UMI counts for each cell were total-counts normalized (no log-normalization). The normalized counts were used to compute an average gene expression level for each marker in each cluster. The cluster-averages for each gene were then standardized to zero-mean and unit-variance for visualization on a common scale.

Differential expression across timepoints was performed separately for each cell type using the “rank_genes_groups” function in scanpy (Wolf et al., 2018). We followed the recommended preprocessing and used default parameters: cells were total-counts normalized and then log transformed with pseudocount 1. A t-test was used to test significance with Benjamini-Hochberg correction for multiple hypotheses. Heatmaps for differentially expressed genes report the degree of enrichment as fold-change over the average expression across timepoints.

### Tamoxifen administration

For temporal cohort analyses (**Figure 4**), pregnant dams were administered 1-3 mg of tamoxifen (stock solution 10 mg/ml in corn oil) via oral gavage, at the corresponding embryonic stage.

### Tissue processing for immunofluorescence staining and FISH

Postnatal animals were transcardially perfused with PBS followed by 4 % paraformaldehyde (PFA) in 120 mM phosphate buffer; their brains were dissected out and post-fixed in 4 % PFA overnight at 4°C. Brains were sectioned into 75-100 µm sections on a vibratome, and either further processed for FISH and/or immunofluorescence staining or stored at 4°C in PBS with 0.05 % sodium azide. Embryonic brains were dissected out in ice-cold PBS and fixed in 4 % PFA overnight at 4°C. The samples were cryoprotected in 30 % sucrose/PBS overnight at 4°C, embedded in O.C.T. compound, frozen on dry ice, and stored at -20°C. Samples were sectioned at 20 µm on a cryostat; sections were either stored at -20 °C or further processed for FISH and/or immunofluorescence staining.

### Immunofluorescence staining

Floating vibratome sections: samples were permeabilized with 0.5 % Triton X-100 in PBS for 1-2 h and blocked in blocking buffer (10 % goat serum, 0.1 % Triton X-100 in PBS) for 1-2 h at room temperature. The sections were then incubated for 24-72 h, at 4°C, with primary antibodies diluted in blocking buffer. The samples were washed three times (10-30 min/wash) with PBS, counterstained with DAPI (4’,6-diamidino-phenylindole) for 45 min (both steps at room temperature), and incubated with secondary antibodies diluted in blocking buffer for 2 h at room temperature or overnight at 4°C. They were then washed (three 10-30 min washes) and mounted on slides with ProLong Gold Antifade Mountant.

Cryosections: Slides were allowed to reach room temperature, and then washed three times with PBS. Sections were permeabilized with 0.5 % Triton X-100 in PBS for 30 min, and blocked with blocking buffer for 1 h at room temperature. Slides were incubated with primary antibodies diluted in blocking buffer overnight, in a humid chamber at 4°C. They were then washed with PBS (three 10-30 min washes), counterstained with DAPI (45 min), and incubated for 1-2 h with secondary antibodies diluted in blocking buffer, at room temperature. Slides were washed (three 10-30 min washes) with PBS and mounted with ProLong Gold Antifade Mountant.

### Fluorescent *in situ* hybridization (FISH)

Embryonic samples at E12 and E14, prepared as outlined above, were submitted to the RNAscope protocol (Advanced Cell Diagnostics), following the manufacturers’ instructions with minor modifications. All RNAscope probes (as listed in the Key Resources table) were purchased from ACD. Additional *in situ* images Allen Developing Mouse Brain Atlas (**Figures 1G,H, S2**) and from the Allen Mouse Brain Atlas (Lein et al., 2007) (**Figure S3B,C**).

### Microscopy and image analysis

Images were acquired using a Leica SP8 laser point scanning confocal microscope. 10x, 25x and 40x objectives were used, and the parameters of image acquisition (speed, resolution, averaging, zoom, z-stack, etc.) were adjusted for each set of samples. Images were further analyzed using ImageJ, both in its native and Fiji distributions, as described below. Brightness and contrast were adjusted as necessary for visualization; the source images were kept unmodified.

## QUANTIFICATION AND STATISTICAL ANALYSIS

### Cell quantification

The CellCounter tool in ImageJ/Fiji was used for all cell quantifications. In the mature septum (**Figures 3 and 4**), all cells positive for the corresponding marker within the dorsal, intermediate and ventral nuclei of the lateral septum (labeled in graphs as LSd, LSi, and LSv, respectively), as well as in the medial septal nucleus (MS) were counted. In **Figure 3D-I**, the rostral, medial and caudal (R/M/C) locations correspond approximately to Bregma +0.75, +0.5 and +0.25, respectively.

Cell morphology types (**Figures 4 and S6**) were determined based on the previous classification by Alonso and Frotscher (Alonso and Frotscher, 1989). Examples are provided in **Figure 4F**. The key descriptive criteria for each neuronal type are:

- Type I neurons have relatively few (3-5) thick and sparsely ramified dendrites, with numerous spines, that form a small, roughly spherical dendritic field surrounding a round or oval soma.
- Type II neurons have thinner and branched dendrites of variable length and orientation and fewer spines, and a round or triangular cell body.
- Type III neurons have thick, spine-dense dendrites that form a bipolar dendritic field from an oval cell body with a characteristic spindle shape.

For quantification of dividing cells in the rostral and caudal portions of the developing septum (**Figure S5**), the septal eminence was identified either by the expression of the tdTomato fluorescent reporter in *Nkx2.1*-expressing cells and their progeny (**Figure S5A,B**), or by immunostaining for NKX2.1 itself (**Figure S5C,D**). In the former case, dividing (late G2/M) cells were identified by pH3 staining; in the latter, cells in M phase (late prophase to late telophase/cytokinesis) were identified from the DAPI counterstaining.

Quantification of RNAscope puncta (**Figures 2 and S3**) was performed using an automated data processing pipeline in MatLab, guided by the SpotsInNucleiBot (https://hms-idac.github.io/MatBots). Each data point corresponds to the average values from the analysis of six fields (dimensions: 100 x 100 µm), located at ventral, intermediate and dorsal positions within the embryonic septum on both hemispheres of a single embryo, analyzed at two different levels along the rostro-caudal axis as indicated. Fields for analyses were obtained from the areas within each sample that had highest levels of expression for the corresponding cell type marker genes (RG – *Nes*; IP – *Ascl1*; NN – *Dcx*), as indicated in **Figure 2C-E**. Values are presented as the density of puncta for each mRNA analyzed normalized to the density of puncta for the corresponding cell type marker.

Cell and RNA puncta numbers were compiled in Microsoft Excel spreadsheets; GraphPad Prism 9 was used to build graphs.

### Statistical analysis

All statistical analyses were performed with GraphPad Prism 9, as detailed in the Figure Legends. All p-values were rounded to ten thousandth, and are presented above each statistical comparison in the corresponding figures; those highlighted in bold are below 0.05, which was considered the cutoff for statistical significance (p-values deemed not statistically significant under this criterion are displayed in regular type).

## AUTHOR CONTRIBUTIONS

Conceptualization: M.T.G., C.C.H.; Formal analysis: S.H., C.W., M.A.A., M.A.N.; Investigation: M.T.G., S.K.S., T.E.L., C.M.R.; Resources: C.C.H.; Data curation: S.H., C.W., M.A.A.; Writing – original draft: M.T.G., C.C.H.; Writing – review & editing: M.T.G., S.K.S., T.E.L., C.M.R., S.H., C.W., C.C.H.; Visualization: M.T.G., S.H., C.W., M.A.A.; Supervision: M.T.G., C.C.H.; Funding acquisition: C.C.H.

## ACKNOWLEDGEMENTS

The authors sincerely thank Alex Ratner and Mandovi Chatterjee at the Single Cell Core at Harvard Medical School for performing the inDrops runs and providing experimental advice; Nicoletta Kessaris, Z Josh Huang, Lucas Cheadle and Michael Greenberg for providing mice; Maria Pazyra and Rosalind Segal for their gift of anti-ZIC antibody; Allon Klein for advice and support; Harwell, Goodrich, Lehtinen, and Segal lab members for discussions and feedback; and Gord Fishell, Lisa Goodrich, Alex Pollen and Matthew Schmitz for their comments on the manuscript.

## FUNDING

M.T.G. was supported in part by the Ellen R. and Melvin J. Gordon Center for the Cure and Treatment of Paralysis. S.K.S. was the recipient of a Boehringer Ingelheim MD Fellowship. T.E.L. is the recipient of a Bill and Melinda Gates Millennium Scholarship from the Gates Foundation; C.M.R. is the recipient of a Gilliam Fellowship for Advanced Study from the Howard Hugues Medical Institute. This work was supported by National Institutes of Health grants R01MH119156 and R01NS102228 and by a Harvard Brain Science Initiative Seed Grant and the Giovanni Armenise Harvard Foundation.

## SUPPLEMENTARY FIGURE LEGENDS

**Figure S1:**
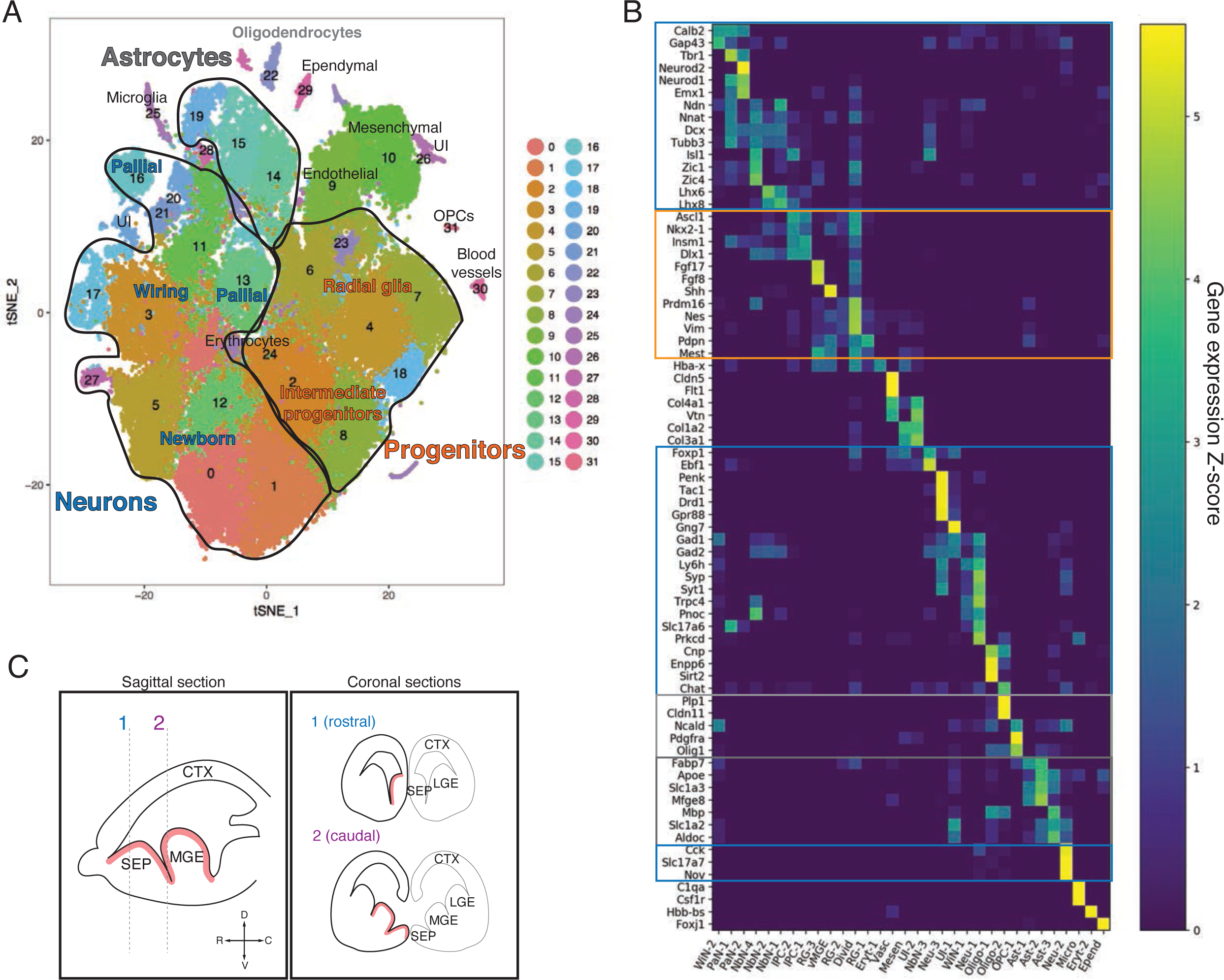
Cluster identity assignment in the scRNA-Seq dataset. **A)** tSNE plot of all cells in the dataset. Assigned identities are written over the corresponding clusters. The three major groups of clusters encircled in black outlines contain progenitor cells (orange text), neurons (blue text) and astrocytes (dark gray text). **B)** Heatmap displaying the relative levels of expression for 73 potential diagnostic genes (rows) in each of the tSNE clusters. Major groups identified in A) are highlighted in the corresponding colors. Putative identities are shown in the x-axis. **C)** Schematic showing the correspondence between sagittal (left) and coronal (right) sections at rostral (1) and caudal (2) positions within the septum (SEP), showing the lateral (LGE) and medial (MGE) ganglionic eminences for reference; the main proliferative area (ventricular zone) is highlighted in red. The compass on left panel indicates the rostro-caudal and dorso- ventral axes). Abbreviations: WiN, wiring neurons; PaN, pallial neurons; NbN, newborn neurons; IPC, intermediate progenitor cells; vMGE, ventral portion of the MGE; RG, radial glia; Divid, dividing cells; Eryt, erythrocytes; Vasc, vasculature; Mesen, mesenchymal cells; UI, unidentified; Neu, neurons; Oligo, oligodendrocytes; OPC, oligodendrocyte progenitor cells; Ast, astrocytes; Micro, microglia; Epend, ependymal cells.

**Figure S2:**
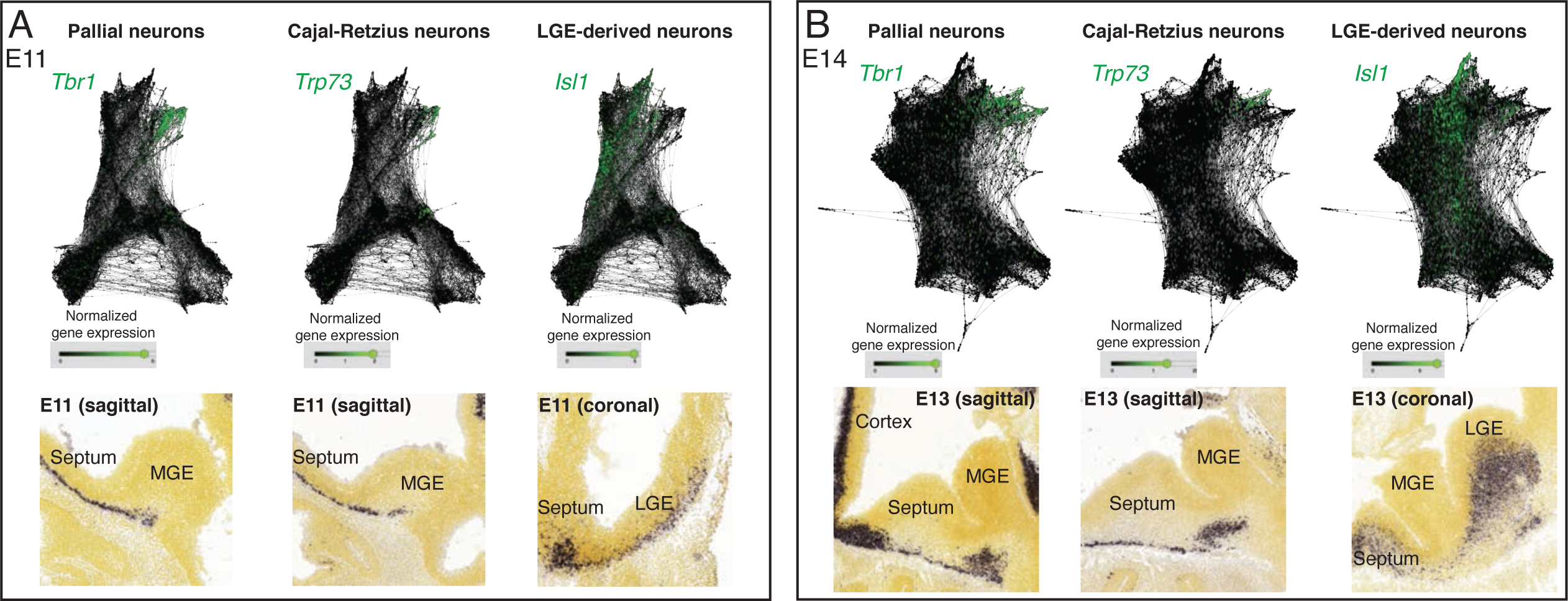
Additional neuronal identities at E11 and E14. **A, B)** Analysis of “protrusions” within the neuronal portion of E11 (A) and E14 (B) SPRING plots shows enrichment in marker genes for newborn pallial neurons (*Tbr1*), Cajal-Retzius cells (*Trp73*) and potentially LGE-like (*Isl1*) newborn neurons present in the septum, analogous across both stages and confirmed by *in situ* hybridization (bottom panels) Image credit (bottom panels): Allen Institute - Allen Developing Mouse Brain Atlas.

**Figure S3:**
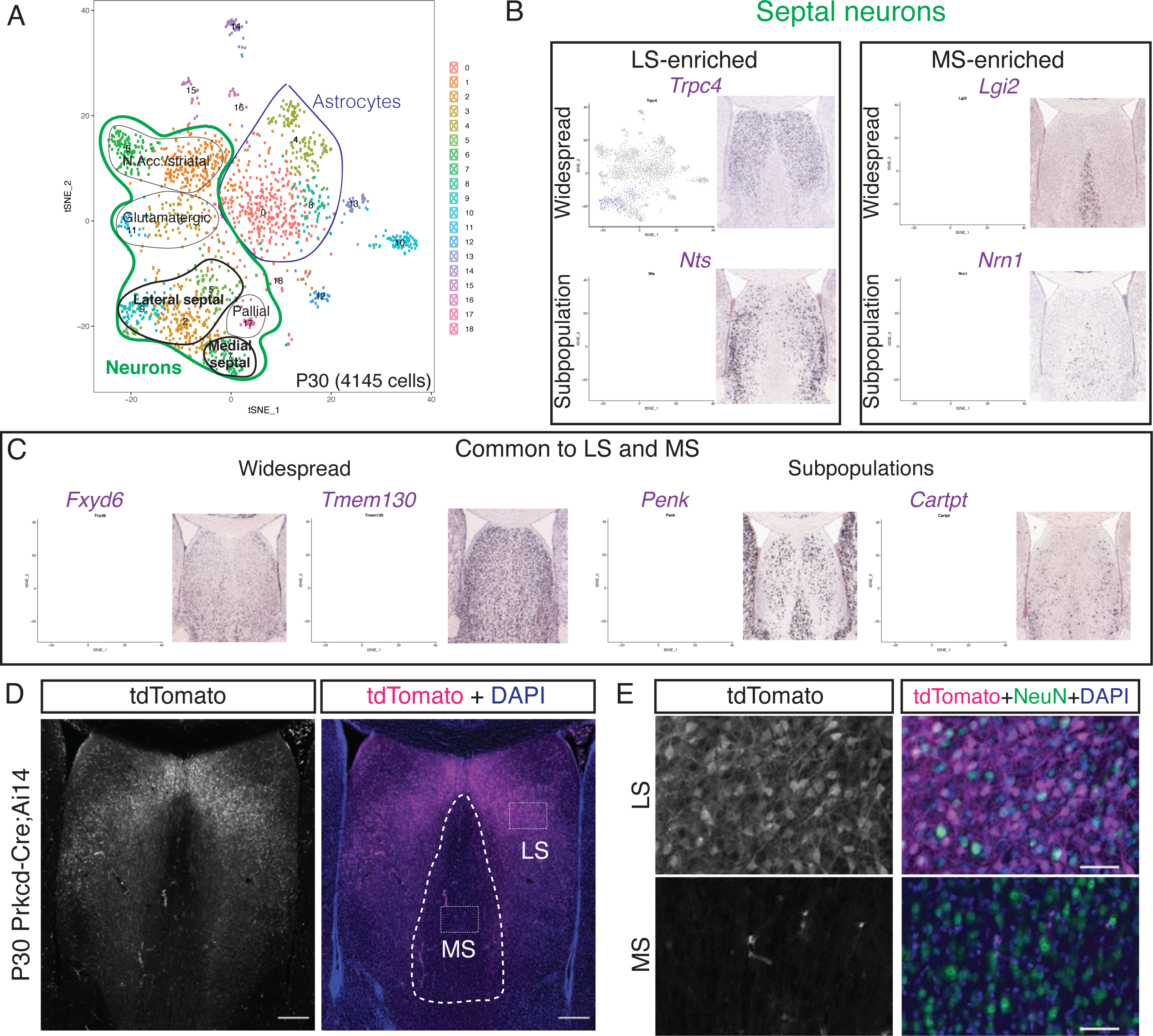
Analysis of P30 dataset for discovery of novel septal neuron markers. **A)** tSNE plot of all P30 cells. Assigned identities are written over the corresponding clusters. Two major cluster groups contain astrocytes (blue), and neurons (green), of which a small proportion are either lateral or medial septal neurons (bold text). **B)** Analysis of gene enrichment in different clusters (feature plots), validated with *in situ* hybridization images from the Allen Brain Atlas, led to the discovery of genes that are enriched in neurons of the LS (left) or MS (right), both following widespread patterns (top) or representing a subpopulation within the corresponding septal nucleus (bottom). Image credit: Allen Mouse Brain Atlas. **C)** Similarly, general septal markers common to LS and MS could be identified, either present in the majority of septal cells (left) or in a subset of them (right). Image credit: Allen Mouse Brain Atlas. **D)** Coronal section through the septum of a P30 Prkcd-Cre;Ai14 mouse, where expression of tdTomato (gray; magenta in merge, counterstained with DAPI in blue), shows that cell labeling is restricted to the lateral septum. Scale bars, 250 µm. **E)** Insets from D), showing overlap of immunostaining for tdTomato (gray; magenta in merge) and NeuN (green in merge) only in the lateral septum.

**Figure S4:**
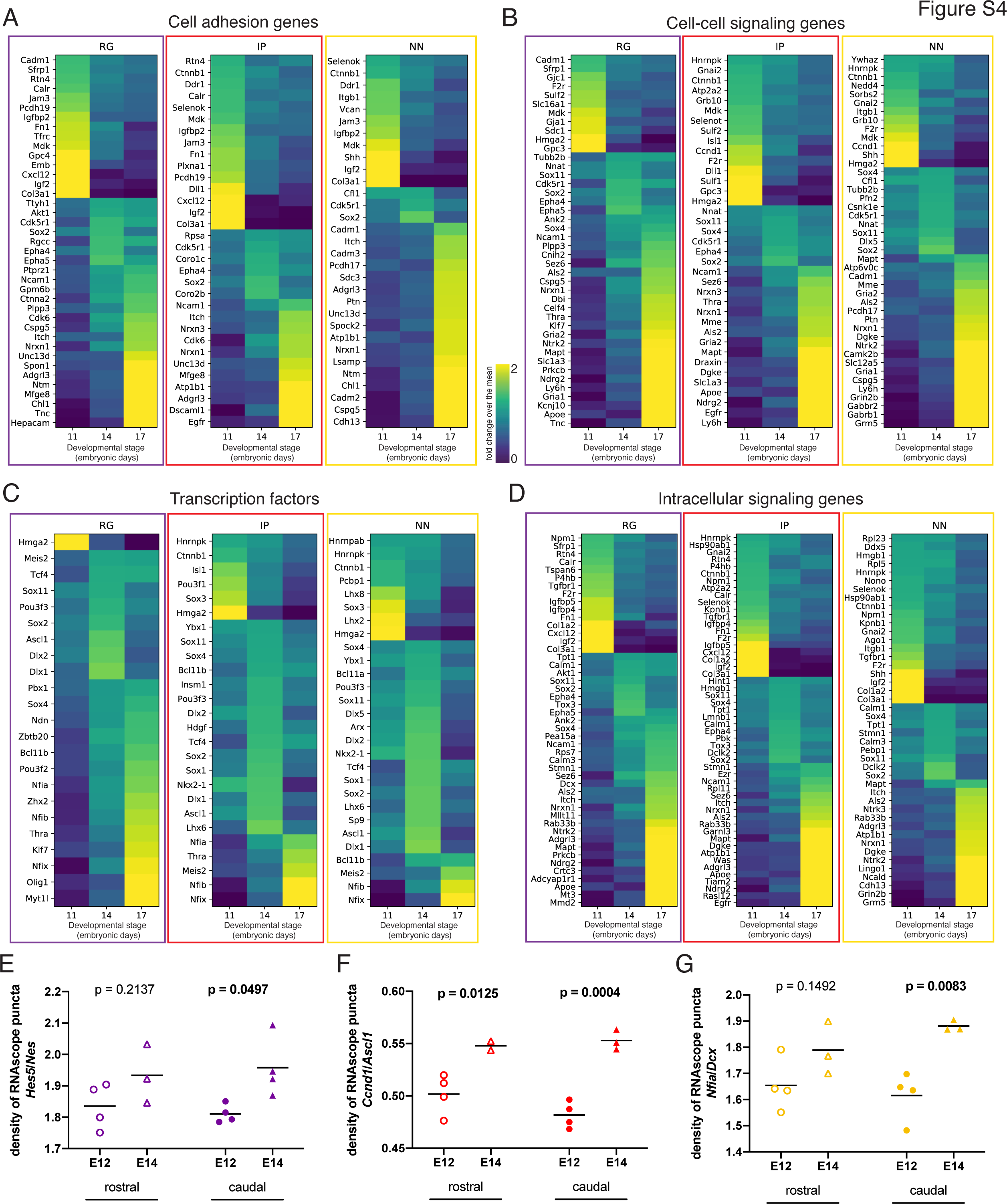
Additional categories and examples of genes differentially enriched across developmental stages. **A-D)** Heatmaps of differentially enriched classes of genes across embryonic stages for the three indicated cell types: radial glia (RG), intermediate progenitors (IP) and newborn neurons (NN), and sorted by biological function category: A) cell adhesion genes; B) cell-cell signaling genes; C) transcription factors; D) intracellular signaling. The color scale applies to panels A-D. **E-F)** Quantification of mRNA puncta density of differentially enriched genes (E, *Hes5*; F, *Ccnd1*; G, *Nfia*), normalized to the density of cell type marker mRNA (E, *Nes*; F, *Ascl1*; G, *Dcx*). Measurements were obtained from the rostral (empty symbols) and caudal (full symbols) portions of the septum at E12 (circles) and E14 (triangles). All data points are represented; black bars represent the mean. Unpaired t-tests were performed; p-values are indicated above the corresponding compared sets of data: those highlighted in bold indicate statistically significant differences (p<0.05).

**Figure S5:**
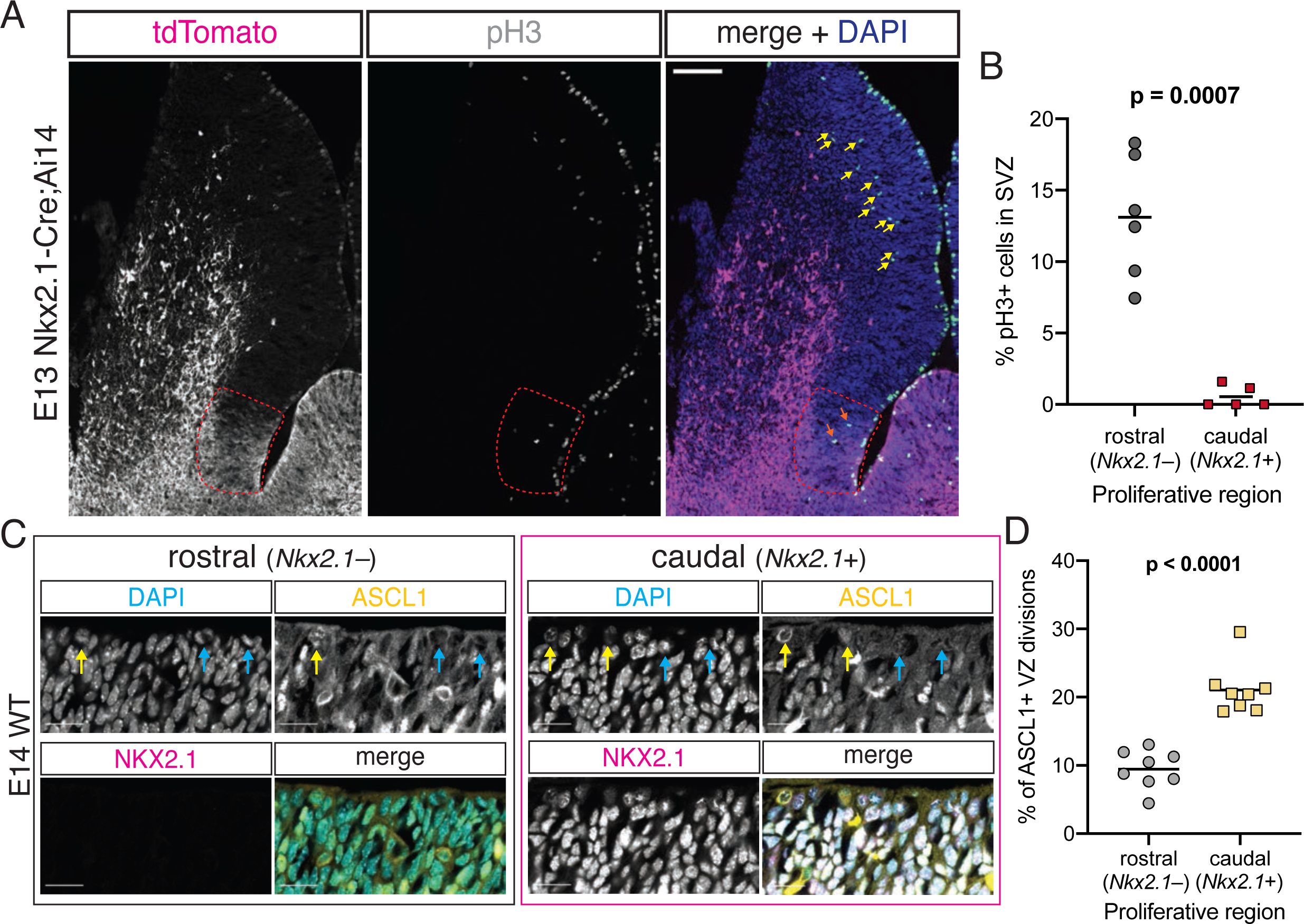
Differences between rostral and caudal septal progenitors. **A)** Closeup of the image displayed in Fig. 3C. Horizontal section of E13 Nkx2.1-Cre;Ai14 mouse brain. Immunofluorescence staining for tdTomato (magenta in merge) and phosphorylated histone 3 (pH3, green in merge) shows difference in the number of subapically dividing cells in the rostral portion of the septum (yellow arrows) compared to the *Nkx2.1*-expressing (red dashed line) septal eminence (orange arrows). Scale bar, 100 µm. **B)** Quantification of the proportion of dividing (pH3+) cells located in the subventricular zone of the rostral (gray dots) and caudal (red squares), i.e. *Nkx2.1*– and *Nkx2.1*+, proliferative regions of the developing septum at E13. **C)** Immunostaining for ASCL1 (yellow in merge) and NKX2.1 (magenta in merge), counterstained with DAPI (blue in merge), in rostral and caudal proliferative regions of the septum of an E14 mouse brain, highlighting ASCL1+ (yellow arrows on DAPI and ASCL1 panels) and ASCL1– (blue arrows on DAPI and ASCL1 panels) dividing cells at the apical surface. Scale bars, 20 µm. **D)** Quantification of the proportion of ventricular surface divisions that are ASCL1+ in the rostral (gray dots) and caudal (yellow squares) proliferative regions of the developing septum at E14. All data points are represented; black bars represent the mean. Unpaired t-tests were performed; the p-values are indicated above the corresponding compared sets of data: bold typeface indicates statistically significant differences (p<0.05).

**Figure S6:**
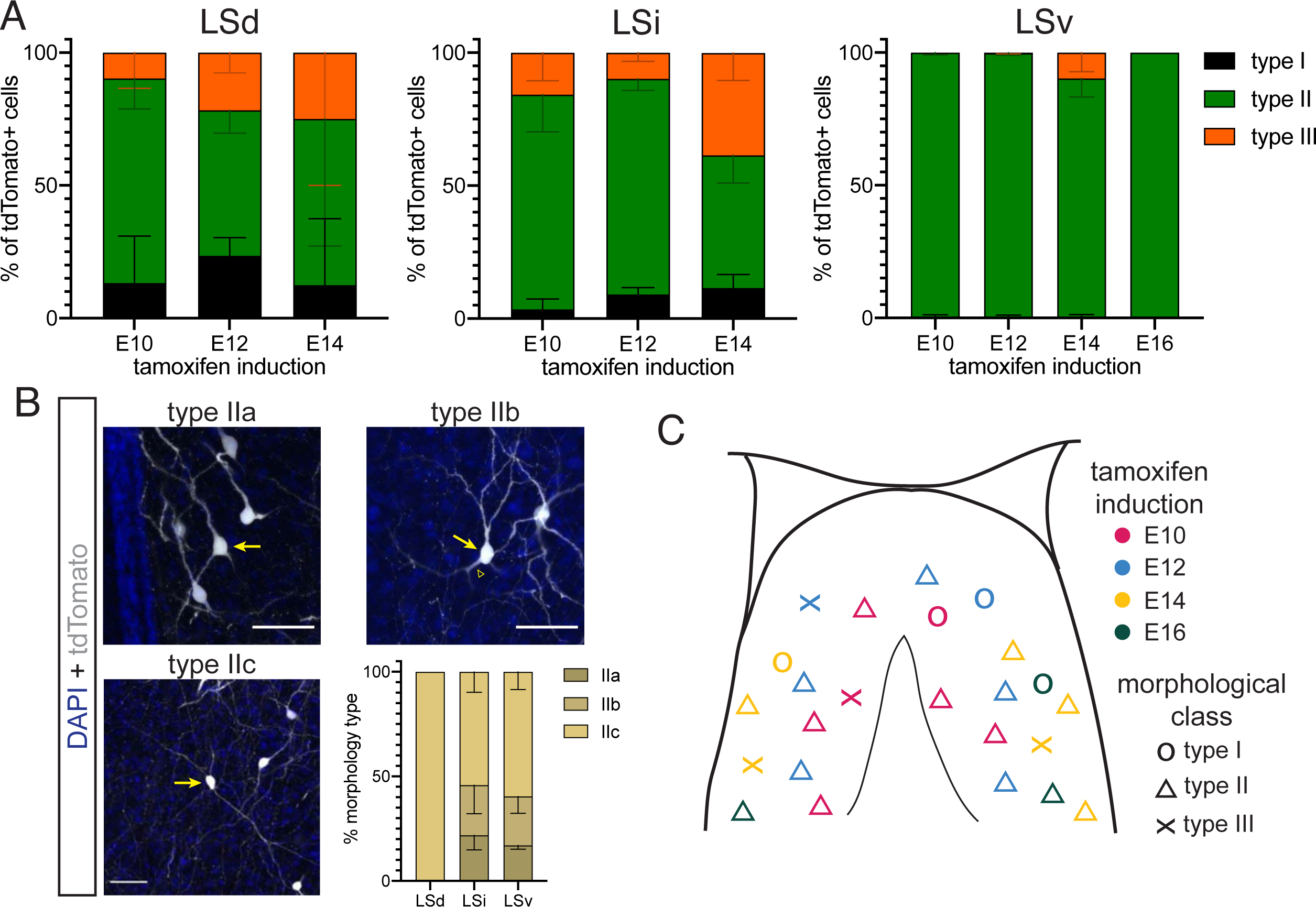
Detailed distribution of morphological cell types across temporal cohorts of septal eminence-derived neurons. **A)** Quantification of the distribution of the three main morphological neuron types in the dorsal (LSd, left), intermediate (LSi, middle) and ventral (LSv, right) nuclei of the lateral septum. Cells in each septal nucleus, represented as a percentage of total classified cells (mean ± s.d.) within each temporal cohort. **B)** Proposed subdivision of morphological neuron type II into three subtypes: IIa (thick dendrites); type IIb (thick initial dendritic segments [arrowhead] that branch out into thinner processes); and type IIc (thin dendrites). Images show cells labeled with tdTomato (gray; counterstained with DAPI, blue); arrows indicate the somata of neurons within each proposed subtype. Quantification of each subtype (as % of total type II cells, mean ± s.d.) within each LS subdivision within the E14 temporal cohort. **C)** Graphic summary of the distribution and morphological types of LS neurons derived from the septal eminence within each temporal cohort.

## Notes

### Competing Interest Statement

The authors have declared no competing interest.

